# Effects of sub-chronic, *in vivo* administration of sigma-1 receptor ligands on platelet and aortic arachidonate cascade in streptozotocin-induced diabetic rats

**DOI:** 10.1101/2022.03.12.484086

**Authors:** Sándor Váczi, Lilla Barna, Krisztián Laczi, Ferenc Tömösi, Gábor Rákhely, Botond Penke, Lívia Fülöp, Tamás Janáky, Mária A. Deli, Zsófia Mezei

**Affiliations:** Department of Pathophysiology, Albert Szent-Györgyi Medical School, University of Szeged, H-6725 Szeged, Hungary; Doctoral School of Theoretical Medicine, University of Szeged, H-6725 Szeged, Hungary; Institute of Biophysics, Biological Research Centre, H-6726 Szeged, Hungary; Doctoral School of Biology, University of Szeged, H-6726 Szeged, Hungary; Department of Biotechnology, University of Szeged, H-6726 Szeged, Hungary; Department of Medical Chemistry, Albert Szent-Györgyi Medical School, University of Szeged, H-6720 Szeged, Hungary; Department of Physiology, Albert Szent-Györgyi Medical School, University of Szeged, H-6720 Szeged, Hungary

**Keywords:** sigma-1 receptor, eicosanoids, platelet, aorta, diabetes, (S)-L1

## Abstract

Diabetes mellitus is a chronic metabolic disorder which induces endothelial dysfunction and platelet activation. Eicosanoids produced from arachidonic acid regulate cellular and vascular functions. Sigma-1 receptor expressed in platelets and endothelial cells can regulate intracellular signalization. Our aim was to examine the influence of sub-chronic, *in vivo* administered sigma-1 receptor ligands (2-(4-morpholino)ethyl-1-phenylcyclohexane-1-carboxylate, PRE-084; S-N-Benzyl-6,7-dimethoxy-1,2,3,4-tetrahydro-1-isoquinolineethanamine, a new compound (S)-L1; and N,N-dipropyl-2-[4-methoxy-3-(2-phenylethoxy)-phenyl]-ethylamine monohydrochloride, NE-100) on the *ex vivo* arachidonic acid metabolism of platelets and aorta in streptozotocin-induced diabetic rats. The serum level of sigma-1 receptor ligands was detected by liquid chromatography-mass spectrometry before the *ex vivo* analysis. Sigma-1 receptor and cyclooxygenase gene expression in platelets were determined by reverse transcription coupled quantitative polymerase chain reaction. The eicosanoid synthesis was examined by using of radiolabeled arachidonic acid substrate and enzyme-linked immunosorbent assay.

In diabetic rats, the sub-chronic, *in vivo* administration of the sigma-1 receptor ligands modified the transcript levels of sigma-1 receptor and cyclooxygenase-1, the concentration of cyclooxygenase in platelets and the eicosanoid synthesis in both platelets and aorta. Sigma-1 receptor ligands, by changing platelet and blood vessel eicosanoid synthesis, may play a role in modulating diabetic complications.

## 1. Introduction

Diabetes mellitus is a chronic, progressive disease characterized by abnormal carbohydrate, lipid and protein metabolism. These abnormalities lead to the formation of glycation end products and accumulation of reactive oxygen species resulting in endothelial dysfunction and platelet activation.^1,2^ The cytokines and eicosanoids synthesized by the impaired endothelium and activated platelets, are involved in the development and progression of atherosclerosis.^3^ Eicosanoids, which regulate cellular and vascular function, are generated from free arachidonic acid (AA) released from phospholipids (PLs) in the cell membrane by phospholipase A_2_ (PLA_2_) in the presence of ionized calcium, by cyclooxygenases (COXs), lipoxygenases (LOXs) and specific synthetases.^4^ The sigma-1 receptor (S1R), which modulates cellular functions,^5–7^ is also expressed on endothelial cells^8^ and platelets.^9^ Beneficial effects of S1R agonists have been already reported in sepsis,^10^ ischemic-reperfusion injury,^11^ stroke^12^ and cardiovascular disease,^13^ as well. Expression of the S1R gene reduces the production of inflammatory cytokines^10^ and protects against the development of diabetic complications.^14^

We aimed to investigate the effects of sigma-1 ligands on platelet and aortic eicosanoid synthesis in streptozotocin-induced (STZ-induced) diabetic rats. We hypothesized that S1R ligands administered sub-chronically *in vivo* (daily for one week) can modulate the *ex vivo* AA metabolism of platelets and aorta when the ligand is not currently present.

For our studies, we chose two well-characterized and one new S1R ligand. PRE-084 (2-(4-morpholino)ethyl 1-phenylcyclohexane-1-carboxylate) was originally identified as a selective agonist of the S1R.^15^ This ligand is able to regulate cytokine production in stroke,^16^ reduce microglial activation in traumatic brain injury, induce protective effects in neurogenic inflammation^17^ and endothelial barrier damage.^18^ NE-100 (N,N-dipropyl-2-[4-methoxy-3-(2-phenylethoxy)-phenyl]-ethylamine monohydrochloride) is an antagonist of the S1R,^19^ which was shown to block sigma-1 agonist (SA4503) induced thoracic aortic vasodilation.^13^ The third ligand we investigated was a recently identified new compound,^20^ (S)-L1 (S-N-benzyl-6,7-dimethoxy-1,2,3,3,4-tetrahydro-1-isoquinolinetanamine), the pharmacological nature of which is still unknown. The serum level of the S1R ligands was determined by liquid chromatography-mass spectrometry (LC-MS) before the *ex vivo* analysis. The eicosanoid synthesis was examined by the use of radiolabeled substrate and enzyme-linked immunosorbent assay (ELISA). The effect of S1R ligands not only depends on the amount and activity of the enzymes, but can also be influenced by the levels of transcripts derived from the genes coding for the S1R and COX, originated from megakaryocytes and stored in platelets. To confirm this process, the S1R *(Sigmar1)* and COX (prostaglandin G/H synthase; *Ptgs*) gene expression in platelets were determined by reverse transcription coupled quantitative polymerase chain reaction (RT-qPCR).

## 2. Materials and Methods

### 2.1. Animals

Animal experiments were performed under the protocol accepted by the Ethical Committee for the Protection of Animals in Research at the University of Szeged, Hungary (Permit No. X./238/2019.). All experiments were carried out in accordance with the Guide for the Care and Use of Laboratory Animals published by the US National Institutes of Health. Male Wistar *(Rattus norvegicus)* rats were used in this study. All animals were maintained in a room, in 12-h dark/12-h light cycles, at constant temperature (23 ± 1°C) with free access to standard laboratory food (SAFE® 132, SAFE, Augy, France) and water ad libitum. As required by guidelines and the ethics permit, we placed enrichment devices (e.g., cylinders, cubes) in the rat cages.

Sample size calculation was performed for the COX metabolism. The estimation was based on the one-way ANOVA hypothesis test. We supposed the effect size to be at least 0.6, statistical power 1-β=0.8 and significance level α=0.05, the calculation resulted in a sample size of 9 per group, 36 animals total. Sample size calculation was performed with G*Power version 3.9.1.7 software.

### 2.2. Diabetic animal model

Diabetes mellitus was induced in Wistar 11-week-old, male rats (n = 36 animals) by a single i.p. injection of 65 mg/bwkg STZ (Sigma, St. Louis, USA) that was dissolved in a freshly prepared 50 mM citrate buffer just before use. After STZ injection, the drinking water of animals was changed for 10% (w/v) sucrose solution for 24 h.^21^ Rats were considered diabetic if the fasting (12 h) peripheral blood glucose concentration was higher than 20 mM 72 h after the injection of STZ (initial serum glucose level). Serum glucose level was monitored by Dcont equipment and Ideal test strips (77 Elektronika Ltd., Budapest, Hungary).

All the animals that received STZ developed diabetes with several fold blood glucose elevation. No mortality was observed and therefore no animals had to be excluded.

### 2.3. Experimental animal groups and their examination

Four weeks later, the diabetic animals were randomly divided into four groups (n= 9 animal/groups) and one of these group was treated with the vehicle of ligands (isotonic saline) and other three groups were treated by the different S1R ligands. Therefore, the generated subgroups were as follows: 1) the control/vehicle-treated, 2) the PRE-084-treated (2-(4-morpholino)ethyl-1-phenylcyclohexane-1-carboxylate; MedChem Express USA), 3) the (S)-L1-treated (S-N-Benzyl-6,7-dimethoxy-1,2,3,4-tetrahydro-1-isoquinolineethanamine, a novel high affinity S1R ligand screened in silico from the in-house compound library of Institute of Pharmaceutical Chemistry, University of Szeged,20 and 4) the NE-100-treated (N,N-dipropyl-2-[4-methoxy-3-(2-phenylethoxy)-phenyl]-ethylamine monohydrochloride; Tocris Bioscience, Bristol, UK) groups, each consisting of 9 rats. 0758-003 is a homologue of (S)-L1 used as the internal standard in LC-MS.

We monitored basal, initial (72 h after STZ administration) and terminal (immediately before *ex vivo/in vitro*) fasting blood glucose levels and body weight of all groups of animals we studied. Serum glucose was determined from blood obtained from rat tails using a D-count blood glucose meter and an Ideal test strip. Before the *ex vivo/in vitro* eicosanoid synthesis studies (terminal phase), platelet counts and serum levels of S1R ligands were determined in all rats from blood taken from the abdominal aorta. After serum S1R ligand determination, the remaining plasma of animals in the same group was randomly divided into 3 pools and serum total cholesterol (colorimetric methods), alanine aminotransferase (ALT, IFCC methods/pyridoxal phosphate activation) and urea (UV kinetic reaction) by COBAS 8000 instrument (Roche, Hungary) were determined from these pooled samples.

The determination of *Sigmar1* and *Ptgs* mRNA levels required 1010 platelets per sample. After testing laboratory parameters and AA metabolism, the remaining blood of one animal no longer contained a sufficient number of platelets for this measurement. To guarantee a suitable platelet count for these tests, we used pooled samples from 3 rats that were selected randomly from the 9-animal group, yielding a total of 3 samples.

### 2.4. In vivo treatment of rats by S1R ligands

#### 2.4.1. Treatment protocols

Four weeks after the STZ injection, the animals were treated with the S1R ligands. All S1R ligand-treated diabetic animals received 3 mg/kg body weight of S1R ligand (PRE-084 or (S)-L1 or NE-100) dissolved in 0.9% sodium chloride, i.p. All diabetic vehicle-treated animals received a 0.9% sodium chloride solution i.p. Treatment in all cases was carried out once a day, for a week. The number of animals/group and the dose of S1R ligands were determined based on a previous publication describing the effects of a S1R ligand applied i.p. in rats.^12^

#### 2.4.2. Determination of serum S1R ligands level

##### Reagents and chemicals

All reagents and chemicals were of analytical or LC–MS grade. Acetonitrile (ACN), methanol (MeOH) and water were obtained from VWR Chemicals (Monroeville, PA, USA). Formic acid (FA) and hydrochloric acid were purchased from Fisher Scientific (Portsmouth, NH, USA). Stock solutions were dissolved in water individually at a final concentration of 1 mg/mL. All standard stock solutions were prepared on ice, divided into 100 μL aliquots, and stored at −80 °C until further use.

##### Preparation of plasma samples for the liquid chromatography-mass spectrometry/mass spectrometry (LC-MS/MS)

To a 100 μL diabetic rat plasma sample 10 μL 0.01 M HCl and according to the analyte’s extraction recovery studies.^22,23^ 330 μL ACN (in (S)-L1 and NE-100) determination or 330 μL MeOH (for the measurement of PRE-084) containing 10 μL 0758-003 internal standard (at 120 nM in (S)-L1 assay and 48 nM for the measurement of PRE-084 and NE-100 were added and the mixture was spun for 60 s. The mixture was allowed to rest for 30 min at −20 °C to support protein precipitation. The supernatant was obtained via centrifugation of the mixture for 15 min at 15,000 × g at 4 °C. The supernatant (340 μL – ACN; 370 μL – MeOH) was transferred to a new tube. After concentration under vacuum (Savant SC 110 A Speed Vac Plus, Savant, USA), the samples were reconstituted in 100 μL starting eluent, vortexed and centrifuged. Finally, 2 μL was injected into the LC–MS/MS system for analysis.

##### The calibration curves for (S)-L1, PRE-084 and NE-100

The rat plasma calibration standards of (S)-L1 (7.81–250 nM), PRE-084, and NE-100 (3.91–125 nM) were prepared by adding the working standard solutions into a pool of drug-free rat plasma.^23^ The sample preparation procedure described above was followed.

##### Analysis of plasma samples by the LC-MS/MS

The quantitative analysis of (S)-L1, PRE-084, and NE-100 was performed after chromatographic separation by using tandem MS. An ACQUITY I-Class UPLC™ liquid chromatography system (Waters, Manchester, UK) comprising Binary Solvent Manager, Sample Manager-FL, and Column Manager connected to a Q Exactive™ Plus Hybrid Quadrupole-Orbitrap Mass Spectrometer (Thermo Fisher Scientific, San Jose, CA, USA) equipped with a heated electrospray ion source (HESI-II) was applied to the analysis. Gradient chromatographic separation was performed at room temperature on a Kinetex EVO C18 column (Phenomenex; 100 Å, 50 mm × 2.1 mm, particle size 2.6 μm) protected by a C18 guard column (Phenomenex, Torrance, CA, USA) by using 0.1 % (v/v) aqueous FA as solvent A and ACN containing 0.1 % (v/v) FA as solvent B.^23^ For quantitative MS analysis of (S)-L1, PRE-084 and NE-100 using MS/MS, the parallel reaction monitoring (PRM) data acquisition mode was selected. To reach the best precursor/product transition for quantitation and maximize sensitivity, the optimal fragmentation conditions and collision energies of each analyte were identified. The mass spectrometer was used in positive mode with the following parameters of the HESI-II source: spray voltage at 3.5 kV, capillary temperature at 253 °C, aux gas heater temperature at 406 °C, sheath gas flow rate at 46 L/h, aux gas flow rate at 11 L/h, sweep gas flow rate at 2 L/h, and S-lens radio frequency (RF) level at 50.0 (source auto-defaults). During data acquisition the transitions of the quantifier and qualifier ions of the analyte and the internal standard were monitored.^23^ A divert valve placed after the analytical column was programmed to switch flow onto the mass spectrometer only when analytes of interest eluted from the column (1.5–3.0 min) to prevent excessive contamination of the ion source and ion optics. The washing procedures of the auto sampler before and after injecting samples were programmed to avoid carryover of analytes. The UHPLC system was controlled using MassLynx 4.1 SCN 901 (Waters). Control of the mass spectrometer, data acquisition, and data processing was conducted using Xcalibur™ 4.1 (Thermo Fisher Scientific).

### 2.5. Separation of platelets

Blood was drawn from the rats 20 hours after the last injection of S1R ligands had been received. Under anesthesia (Euthasol®/pentobarbital-Na/ 30 mg/bwkg i.p.), blood was drawn from the abdominal aorta of rats with a thick needle and was diluted (1:2) with phosphate buffer (pH 7.4) containing ethylene diamine tetra acetic acid (EDTA, 5.8 mM) and glucose (5.55 mM). Platelets were separated by differential centrifugation.^24,25^ Platelet-rich plasma was collected after the whole blood had been centrifuged at 200 g, for 10 min, at room temperature. Platelets were sedimented from the supernatant by centrifugation at 2000 g, for 10 min. The pellet was contaminated with red blood cells; therefore, erythrocytes were lysed with hypoosmotic ammonium chloride (0.83%, 9 parts) containing EDTA (0.02%, 1 part) over 15 min. The platelets were then washed twice with phosphate buffer pH 7.4 containing 5.8 mM EDTA and 5.55 mM glucose, and centrifuged at 2000 g, for 10 min, at room temperature to remove the ammonium chloride and the erythrocyte residues/debris. The absolute platelet count was determined before the second centrifugation. After the last centrifugation, the platelets were resuspended (2.5×10^8^ platelets/mL) in serum-free Medium 199 tissue culture (Sigma, St. Louis, MO).

### 2.6. Determination of Sigmar1, Ptgs1 and Ptgs2 gene expression by RT-qPCR

Rat platelet samples were homogenized in TRI Reagent (Sigma-Aldrich, USA), then RNA from each sample was transcribed to complementary DNA by High Capacity cDNA Reverse Transcription Kit (Applied Biosystems, USA) according to the manufacturer’s protocol based on random priming. The RT-qPCR was performed with TaqMan Gene Expression Master Mix (Life Technologies, USA) in a CFX96 Touch Real-Time PCR Detection System (Bio-Rad Laboratories, USA). Inventoried TaqMan Gene Expression Assays (Life Technologies, USA) were the following: *Sigmar1* - Rn00578590_m1; *Ptgs1* - Rn00566881_m1; *Ptgs2* - Rn01483828_m1; glyceraldehyde 3-phosphate dehydrogenase (Gapdh) - Rn01749022_g1. After heat activation at 95 °C for 3 min the cycling conditions were the following: denaturation for 30 s at 95 °C, amplification for 30 s at 60 °C (40 cycles). qPCR data were analyzed by Bio-Rad CFX Maestro software (Bio-Rad Laboratories, USA). In all samples the transcript level of a gene was normalized to an endogenous control gene (Gapdh, ΔCt = Ct gene – Ct Gapdh). Then ΔΔCt was calculated in comparison to the relative expression of the target genes in vehicle-treated control groups. Fold changes were calculated using the 2^−ΔΔCt^ formula.

### 2.7. Isolation of aorta

Under anesthesia (Euthasol®/pentobarbital-Na/ 30 mg/kg body weight i.p.), after blood collection, the abdominal aorta of the rat was isolated from the branch of the iliac artery to the diaphragm. At 4°C temperature, the connective tissue was removed from the aorta, which was sliced into 1–2 mm thick rings with care to not damage the endothelium.^26^

### 2.8. Examination of AA metabolism

#### 2.8.1. Analysis of eicosanoid synthesis by isotope-labeled AA

An *ex vivo* examination of the eicosanoid synthesis of platelets was carried out in Medium 199, which consists of mineral salts, nucleotides, amino acids, vitamins, carbohydrates, and Ca^2+^ in the same amount as the extracellular concentration but without fibrinogen or other plasma proteins. The diabetic (*in vivo* vehicle-treated and PRE-084-treated, or (S)-L1-treated, or NE-100-treated) rat platelets (2.5×10^8^ platelets/mL in each sample) were pre-incubated at 37°C for 3 min, while the rat abdominal aortic rings (15 mg wet mass/mL) were pre-incubated for 10 min in Medium 199 (Merck). The examined enzyme reaction started after pipetting the tracer substrate 1-^14^C-AA (3.7 kBq, 0.172 nM in each sample; American Radiolabeled Chemicals, Inc., St Louis, MO 63146 USA) into the incubation mixture. The enzyme reaction was stopped by bringing the pH to 3 with formic acid, in case of the platelet samples after 13 minutes of incubation, and in case of the aortic samples after 30 minutes of incubation. The samples were extracted with ethyl acetate and the organic phases were evaporated. The residues were reconstituted in ethyl acetate and quantitatively applied to silica gel G thin-layer plates (Kieselgel G 60/DC-Fertigplatten, Merck, Art. 5721). The plates were developed to a distance of 16 cm in the organic phase of ethyl acetate:acetic acid:2,2,4-trimethylpentane:water (110:20:30:100) by using overpressure thin-layer chromatography (Chrompress 25, Labor MIM, Hungary).^26–29^ A semi-quantitative analysis of labeled AA metabolites was performed by a BIOSCAN AR-2000 Imaging Scanner (Eckert & Ziegler Radiopharma, Berlin, Germany) using Win-Scan 2D Imaging Software. The radio-labeled products of AA were identified with unlabeled authentic standards.^24^ Assuming that the exogenously administered labeled AA, as a tracer, is converted in the same way as the endogenous source, our method allows measurement of the relative amounts of various prostanoids.^26^

#### 2.8.2. Analysis of eicosanoid synthesis by ELISA

The course of this procedure is the same as the examination of radioactive AA metabolism (see above), but in this case, the radioactive-labeled substrate is not added to the incubation mixture. After incubation, the incubation mixture, which contained either platelets or aortic rings, was placed at −80°C and stored at this temperature until the assay was performed. Before performing the ELISA test, the samples were lyophilized and the lyophilizates were re-suspended in the 310 μL 0.9% NaCl solution as the original incubation medium. The prepared samples were centrifuged at 3000 rpm, for 10 min to remove cell debris. Subsequently, the amount of TxB_2_ (sensitivity: 5.48 ng/L; assay range: 5–1800 ng/L), COX-1(sensitivity: 0.285 ng/mL, assay range: 0.3-60 ng/mL), and COX-2 (sensitivity: 0.446 ng/mL, assay range: 0.5-150 ng/mL) was determined in the diabetic rat platelets and aorta by ELISA kits (Shanghai Sunred Biological Technology Co. Ltd, PRC), in compliance with the protocols attached to the kits. The optical absorbance of the samples was measured at 450 nm using a STAT FAX 2100 ELISA plate reader (Awareness Technology, Inc. 34991 Palm City, FL, USA). Based on the calibration curves, the respective concentrations of the examined metabolites and enzymes were determined using the SPSS 22.0 software.

### 2.9. Statistical analysis

The physical and laboratory parameters of animals, the eicosanoid synthesis results of ELISA and the labeled AA substrate were assessed by one-way analysis of variance (ANOVA) followed by Bonferroni’s post hoc test. The results of RT-qPCR tests were assessed by one-way analysis of variance followed by Dunnett’s post hoc test. A difference at a level p < 0.05 was considered statistically significant. The results are expressed as means ± S.D. The statistical analysis was performed by SPSS version 22.0 (IBM Corp. Released 2013. IBM SPSS Statistics for Windows, Version 22.0. Armonk, NY: IBM Corp.)

## 3. Results

### 3.1 Physical and laboratory parameters of animals

#### 3.1.1. STZ-induced diabetic model

All STZ-treated animals had fasting blood glucose levels several times the age- and sex-matched reference range published by Charles River Laboratories,^30^ i.e. hyperglycemia. In addition to hyperglycemia, the development of diabetes was confirmed by the fact that the animals' body weight gain was significantly below the sex- and age-matched physiological rat body weight published by Charles River Laboratories.^30^ The terminal blood glucose levels, body weights and platelet counts of diabetic animals receiving different treatments (vehicle, PRE-084, (S)-L1, NE-100) did not show significant differences *(Table 1)*.

**Table 1.** Body weight, fasting serum glucose levels and platelet number of diabetic rats.

|  | *Basal* | *Initial* | *Terminal* | *Basal* | *Initial* | *Terminal* | *Terminal* |
| --- | --- | --- | --- | --- | --- | --- | --- |
| **Vehicle** | 314±25 | 319±23 | 340±33 | 6.5±0.3 | 29±2.9*** | 32±2.7*** | 522±79 |
| **PRE-084** | 317±36 | 322±36 | 347±42 | 6.4±0.3 | 29±3.1*** | 33±2.3*** | 591±132 |
| **(S)-L1** | 326±13 | 331±13 | 369±27 | 6.5±0.4 | 28±4.1*** | 32±3.8*** | 507±86 |
| **NE-100** | 315±15 | 320±16 | 333±18 | 6.5±0.3 | 30±2.4*** | 32±2.4*** | 635±67 |

Data is shown as mean ± SD; n= 9-9 animals/groups; One-way ANOVA, Bonferroni test, \*\*\*p < 0.001 initial or terminal serum glucose levels were compared to the basal serum glucose levels.

At the end of the *in vivo* study from the pooled blood samples the levels of total cholesterol, alanine aminotransferase (ALT) and blood urea nitrogen (BUN) were determined *(Table 2)*.

**Table 2.** Laboratory parameters of diabetic rats at the end of the *in vivo* study (terminal phase).

| **RATS** | **Total cholesterol** | **ALT (U/L)** | **BUN (mM)** |
| --- | --- | --- | --- |
| **Vehicle** | 2.57±0.078 | 82.0±0.57 | 11.03±0.32 |
| **PRE-084** | 2.78±0.012 | 112.5±7.50*** | 8.97±0.31** |
| **(S)-L1** | 3.15±0.57 | 136.33±10.02*** | 9.97±1.05 |
| **NE-100** | 2.42±0.035 | 145.30±3.51*** | 10.33±0.55 |

Data is shown as mean ± SD; n=3-3 pooled samples/group from 9-9 animals/groups; One-way ANOVA, Bonferroni test, \*\*p < 0.03, \*\*\*p < 0.01 compared to the vehicle treated group; ALT: Alanine Aminotransferase; BUN: blood urea nitrogen

The serum total cholesterol, ALT and urea levels of STZ-induced diabetic animals treated with vehicle were higher than those of healthy, sex- and age-matched rats (serum cholesterol: 1.5±0.34 mM; ALT: 28±7 U/L; BUN: 2.85±0.48 mM).^30^ None of the S1R ligands that we tested significantly altered the increase in serum total cholesterol levels induced by STZ treatment. However, all three S1R ligands further enhanced the increase in ALT levels observed in vehicle-treated diabetic rats. The smallest increase in ALT was detected when PRE-084 was used. However, the STZ-induced increase in serum urea levels was only attenuated by PRE-084 *(Table 2)*.

#### 3.1.2. Concentration of S1R ligands in rat plasma

In our preliminary experiments we demonstrated that intraperitoneal (i.p.) administered Sigma-1 ligands appear in the circulation within one hour, but after 20 hours their plasma levels are at the limit of detection.^23^

Plasma concentrations of S1R ligands were determined in all of the diabetic rats 20 hours after the last ligand injection, immediately before the *ex vivo* studies. Plasma concentrations of S1R ligands were determined in all of the diabetic rats 20 hours after the last ligand injection, immediately before the *ex vivo* studies. In the diabetic rats, the elimination of PRE-084 from the circulation was the fastest of all the Sigma-1 ligands we tested, while that of (S)-L1 was the slowest *(Table 3)*.

**Table 3.** Concentration of S1R ligands in the plasma of diabetic rats.

| ANALYTE | CONCENTRATION (nM) |
| --- | --- |
| **PRE-084** | < LOQ |
| **(S)-L1** | 10.99±1.06 |
| **NE-100** | 5.31±3.14 |

Samples were taken 20 hours after the last dose (3 mg/bwkg/day for 7 days). Data is shown as mean ± SD; n = 9-9 rats; LOQ: limit of quantification.

### 3.2. Effects of S1R ligands on the level of Sigmar1 mRNA in rat platelets

The mRNA levels of *Sigmar1* were detected in rat platelet samples using RT-qPCR *(Fig. 1)*.

**Fig 1.**
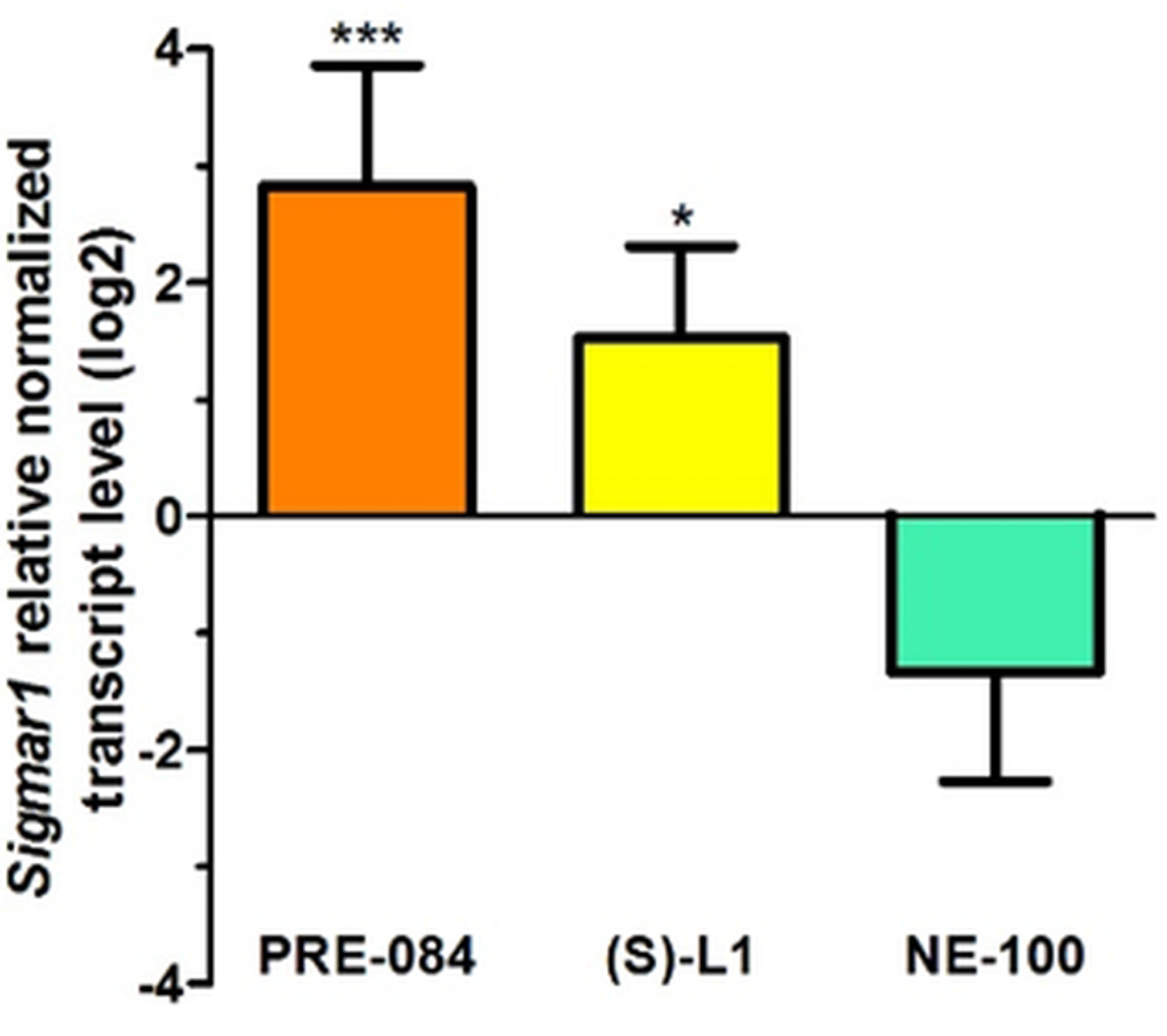
Relative normalized transcript levels of *Sigmar1* in rat platelet samples from diabetic rats treated with S1R ligands compared to diabetic vehicle-treated group.

Rats were treated with 3 mg/kg/day PRE-084, (S)-L1 or NE-100 for 7 days. The vehicle-treated group received saline solution. The transcript levels of *Sigmar1* in rat platelet samples were first normalized to their own endogenous control mRNA levels (*Gapdh*) and then to the similarly normalized mRNA levels in diabetic vehicle-treated animal groups. Fold changes were calculated using the 2^−ΔΔCt^ formula. Mean ± SD, n=3 samples pooled from 9 rats, one-way ANOVA, Dunnett test, *p < 0.05, ***p < 0.001 compared to the diabetic vehicle-treated groups.

The *Sigmar1* mRNA showed a low concentration level in diabetic rat platelets. Although Ct values were high in most of the cases, we considered them reliable, because the RT-qPCR was performed with TaqMan assays. In diabetic rats, the daily administration of PRE-084 and (S)-L1 for one week, elevated *Sigmar1* mRNA levels as compared to samples from vehicle-treated diabetic rats. The *Sigmar1* mRNA level was lower in the NE-100-treated diabetic group compared to those in PRE-084 and (S)-L1 treated samples, but did not differ significantly from the vehicle-treated diabetic group *(Fig. 1)*.

### 3.3. Effects of S1R ligands on the level of Ptgs1 mRNA in rat diabetic platelets

In diabetic rats PRE-084 treatment did not change *Ptgs1* mRNA levels as compared to samples from vehicle-treated diabetic rats. The *Ptgs1* mRNA level was lower in the (S)-L1 group than in the PRE-084 group of diabetic rats, but did not differ significantly from the vehicle-treated diabetic group. NE-100 treatment in diabetic rats significantly decreased the platelet *Ptgs1* level as compared to the vehicle-treated diabetic group *(Fig. 2)*.

**Fig 2.**
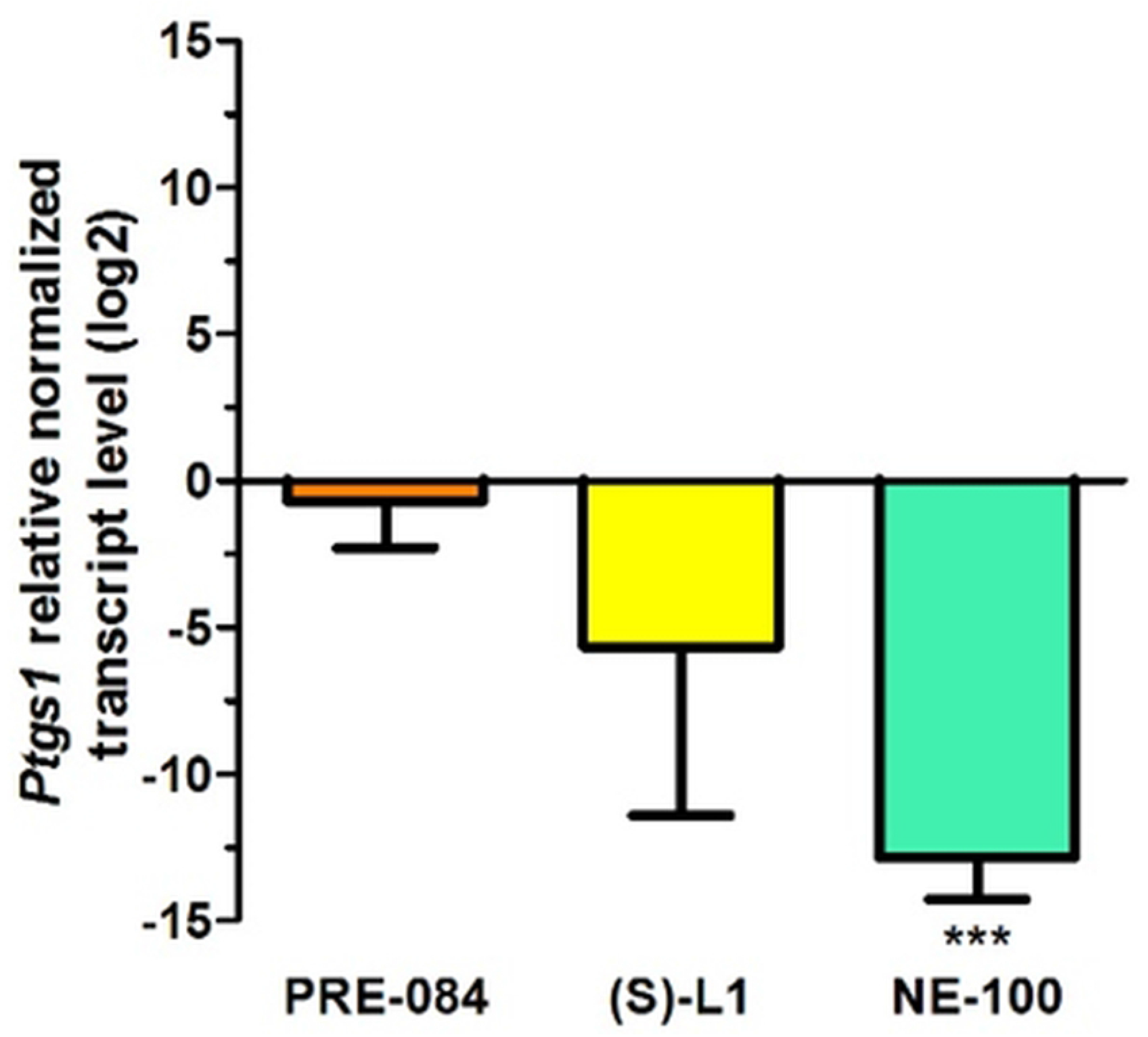
Relative normalized transcript levels of *Ptgs1* in rat platelet samples from diabetic rats treated with S1R ligands compared to diabetic vehicle-treated group.

Rats were treated with 3 mg/kg/day PRE-084, (S)-L1 or NE-100 for 7 days. The vehicle-treated group received only saline solution. The transcript levels of *Ptgs1* in the rat platelet samples were first normalized to their own endogenous control mRNA levels (*Gapdh*) and then to the similarly normalized mRNA levels in diabetic vehicle-treated animal groups. Fold changes were calculated using the 2^−ΔΔCt^ formula. Mean ± SD, n=3 samples pooled from 9 rats, one-way ANOVA, Dunnett test, ***p < 0.001 compared to the diabetic vehicle-treated groups.

Interestingly, the effect of (S)-L1 was different in the case of *Sigmar1* and *Ptgs1*mRNA levels: an effect similar to the S1R agonist PRE-084 was observed for *Sigmar1* mRNA level (Fig. 1), while a trend similar to the S1R antagonist NE-100 was seen for *Ptgs1* mRNA concentration in rat platelets *(Fig. 2)*.

We measured but could not detect *Ptgs2* mRNA by RT-qPCR in platelet samples using 40 cycles.

### 3.4. Analysis of ex vivo AA metabolism in diabetic rats

#### 3.4.1. Effects of S1R ligands on the *ex vivo* eicosanoid synthesis of platelets

##### Application of radioactive AA substrate to study platelet eicosanoid synthesis

In diabetic rat platelets, the total amount of the COX metabolites (sum of 6-keto prostaglandin F_1α_/6-k-PGF_1α_, that is a stable metabolite of prostacyclin; prostaglandin F_2α_/PGF_2α_; prostaglandin E_2_/PGE_2_; prostaglandin D_2_/PGD_2_; thromboxane B_2_/TxB_2_, which is a stable metabolite of TxA_2_, and 12-L-hydroxy-5,8,10-heptadecatrienoic acid/12-HHT) was significantly reduced by PRE-084 and (S)-L1, while it was not affected by NE-100, compared to the vehicle-treated diabetic rat platelets. The total amount of the LOX metabolites as well as the total amount of the AA metabolites (COX+LOX) in diabetic platelets significantly decreased in the NE-100 treated group, compared to the vehicle-treated diabetic samples. The COX/LOX ratio was significantly decreased in the PRE-084 or (S)-L1 treated groups, while it was not modified by NE-100 compared to the vehicle-treated diabetic samples *(Table 4)*.

**Table 4.** Arachidonic acid metabolism in diabetic rat platelets.

| RATS | COX (cpm) | LOX (cpm) | COX+LOX (cpm) | COX/LOX |
| --- | --- | --- | --- | --- |
| **Vehicle** | 1599±164 | 3320±268 | 4919±384 | 0.48±0.02 |
| **PRE-084** | 1408±85*** | 3421±126 | 4835±111 | 0.41±0.01*** |
| **(S)-L1** | 1414±147* | 3434±181 | 4853±250 | 0.41±0.02** |
| **NE-100** | 1576±83 | 2988±204** | 4557±249* | 0.53±0.01 |

**cpm:** count per minute; **COX:** total amount of cyclooxygenase metabolites synthesized from AA in platelets; **LOX:** total amount of lipoxygenase metabolites synthesized from AA in platelets; All data are represented as mean ± SD. n=9–9 rats/group. One-way ANOVA, Bonferroni test, \**p* < 0.05, \*\**p* < 0.03, \*\*\**p* < 0.01 compared to the diabetic vehicle-treated groups.

PRE-084 and (S)-L1 ligands induced the elevation of TxB_2_ (stable metabolite of TxA_2_, the main vasoconstrictor and platelet aggregator product) and the reduction of PGD_2_ (vasodilator and platelet anti-aggregator product) synthesis in the diabetic platelets, compared to the vehicle-treated diabetic platelets *(Fig. 3)*.

**Fig 3.**
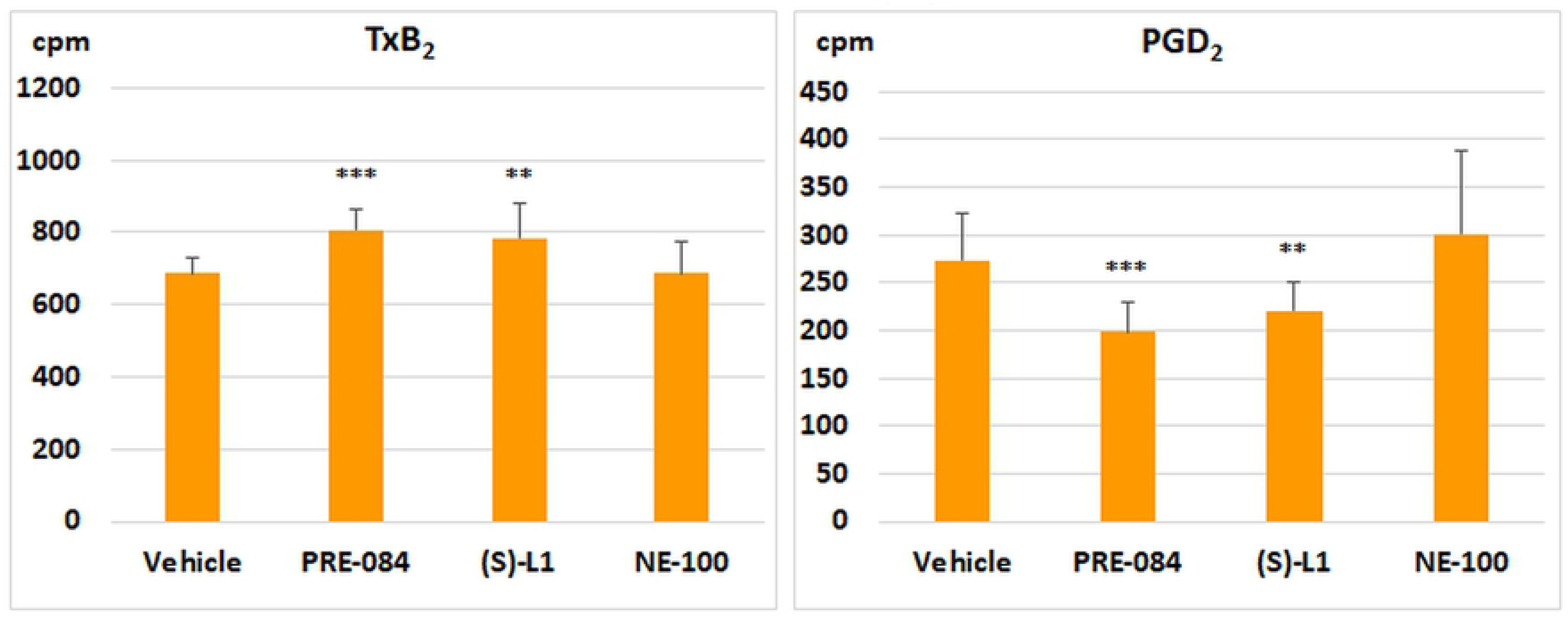
The effects of S1R ligands on the different COX metabolites in diabetic rat platelets.

Y axis represents the isotope activity in count per minute (cpm); All data are represented as mean ± SD. n=9−9 rats/group. One-way ANOVA, Bonferroni test, \*\**p* < 0.03, \*\*\**p* < 0.01 compared to the diabetic vehicle-treated groups.

The amount of vasoconstrictor and platelet aggregator COX metabolites (CON: the sum of PGF_2_α and TxB_2_) was increased in diabetic platelets by PRE-084 ligand. However, the synthesis of vasodilator and anti-aggregator COX metabolites (DIL: sum of PGE_2_ and PGD_2_) was reduced not only by PRE-084 but also by (S)-L1 ligand, compared to the vehicle-treated diabetic group. In the platelets of diabetic rats, an increase in CON/DIL ratio was observed in both PRE-084 and (S)-L1 treated groups, compared to the diabetic, vehicle-treated rat platelets *(Table 5)*.

**Table 5.** Vasoconstrictor, platelet aggregator (CON) and vasodilator, platelet anti-aggregator (DIL) COX metabolites of diabetic rat platelets.

| RATS | CON (cpm) | DIL (cpm) | CON/DIL |
| --- | --- | --- | --- |
| **Vehicle** | 786±51 | 472±68 | 1.68±0.22 |
| **PRE-084** | 875±63*** | 311±44*** | 2.84±0.41*** |
| **(S)-L1** | 836±103 | 331±28*** | 2.56±0.48*** |
| **NE-100** | 809±85 | 514±88 | 1.57±0.31 |

**cpm:** count per minute; **CON:** the sum of $\text{PGF}_2\alpha$ and $\text{TxB}_2$ ; **DIL:** the sum of $\text{PGE}_2$ and $\text{PGD}_2$ ; All data are represented as mean ± SD. n=9–9 rats/group. One-way ANOVA, Bonferroni test, \*\*\* $p < 0.01$ compared to the diabetic vehicle-treated groups.

##### COX enzymes levels in diabetic rat platelets determinate by ELISA

In diabetic rat platelets, the concentration of the COX-1, that is a constitutive isoform of COX enzyme, was increased by PRE-084 and (S)-L1 ligands. However, the amount of COX-2, that is an inducible isoform of COX enzyme, was elevated by all of the examined S1R ligands (PRE-084, (S)-L1, NE-100). Despite the fact that all three ligands we tested increased the total amount of COX enzymes (COX-1 + COX-2), only NE-100 induced a shift in the COX-1/COX-2 ratio towards inducible COX-2 *(Table 6)*.

**Table 6.**
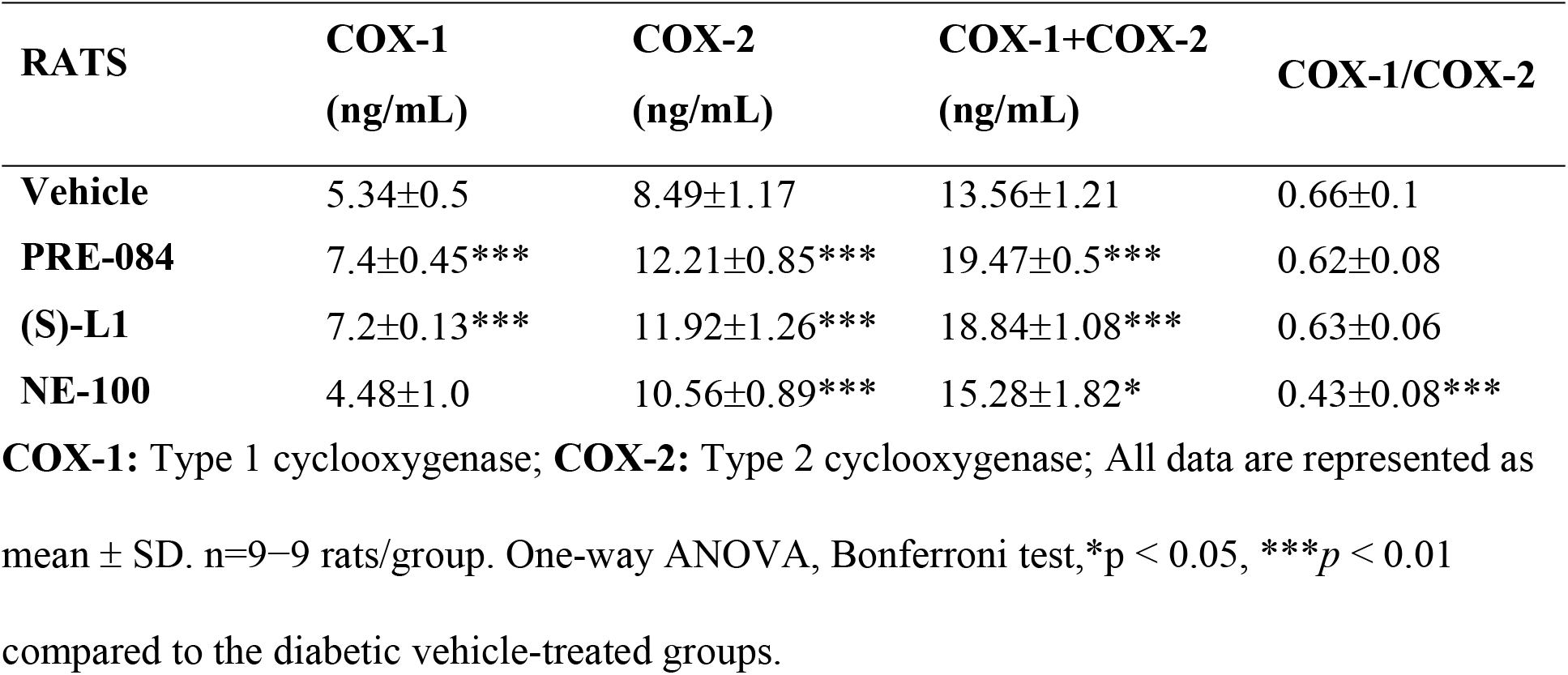
Effect of S1R ligands on the amount of COX-1 and COX-2 enzymes determined by ELISA, in diabetic rat platelets.

| RATS | COX-1<br>(ng/mL) | COX-2<br>(ng/mL) | COX-1+COX-2<br>(ng/mL) | COX-1/COX-2 |
| --- | --- | --- | --- | --- |
| **Vehicle** | 5.34±0.5 | 8.49±1.17 | 13.56±1.21 | 0.66±0.1 |
| **PRE-084** | 7.4±0.45*** | 12.21±0.85*** | 19.47±0.5*** | 0.62±0.08 |
| **(S)-L1** | 7.2±0.13*** | 11.92±1.26*** | 18.84±1.08*** | 0.63±0.06 |
| **NE-100** | 4.48±1.0 | 10.56±0.89*** | 15.28±1.82* | 0.43±0.08*** |

**COX-1:** Type 1 cyclooxygenase; **COX-2:** Type 2 cyclooxygenase; All data are represented as mean ± SD. n=9–9 rats/group. One-way ANOVA, Bonferroni test, \* $p < 0.05$ , \*\*\* $p < 0.01$ compared to the diabetic vehicle-treated groups.

#### 3.4.2. Effects of S1R ligands on the *ex vivo* eicosanoid synthesis of the diabetic rat abdominal aorta

##### Application of the radioactive AA substrate for the study of aortic eicosanoid synthesis

The synthesis of COX and LOX metabolites was also increased in the diabetic rat abdominal aorta in the NE-100 treated group, compared to the vehicle-treated diabetic one. PRE-084 and (S)-L1 administration reduced the production of LOX metabolites, without modifying the COX pathway in the diabetic aorta compared to the vehicle-treated diabetic aorta. The amount of AA metabolites (COX+LOX) in the diabetic rat abdominal aorta was significantly reduced in the (S)-L1 treated group, but was increased in the NE-100 treated group, compared to the vehicle-treated diabetic animals. The COX/LOX metabolite ratio in the diabetic rat aorta significantly increased in the PRE-084 treated group, compared to the vehicle-treated diabetic rats *(Table 7)*.

**Table 7.** Arachidonic acid metabolism in diabetic rat abdominal aorta.

| RATS | COX (cpm) | LOX (cpm) | COX+LOX (cpm) | COX/LOX |
| --- | --- | --- | --- | --- |
| **Vehicle** | 815±201 | 555±94 | 1369±281 | 1.46±0.2 |
| **PRE-084** | 761±187 | 370±55*** | 1132±183 | 2.13±0.7* |
| **(S)-L1** | 658±238 | 373±74*** | 1031±300* | 1.73±0.4 |
| **NE-100** | 1060±102*** | 800±64*** | 1860±131*** | 1.33±0.1 |

**cpm:** count per minute; **COX:** total amount of cyclooxygenase metabolites synthesized from AA in abdominal aorta; **LOX:** total amount of lipoxygenase metabolites synthesized from AA in abdominal aorta; All data are represented as mean ± SD. n=9–9 rats/group. One-way ANOVA, Bonferroni test, \*p < 0.05, \*\*\*p < 0.01 compared to the diabetic vehicle-treated groups.

(S)-L1 and NE-100 ligands induced the reduction of TxB_2_ synthesis and the elevation of 6-k-PGF_1α_ synthesis in the diabetic aorta, compared to the vehicle-treated diabetic ones *(Fig. 4)*.

**Fig 4.**
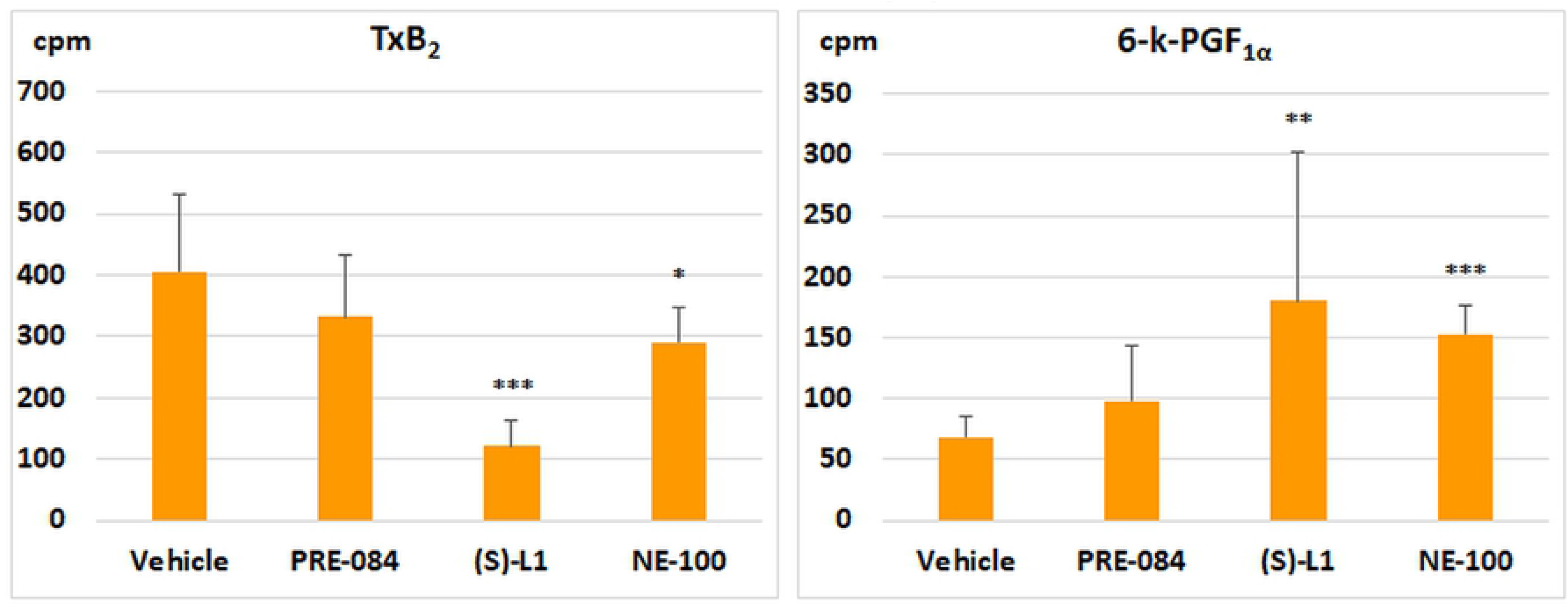
The effects of S1R ligands on the different COX metabolites in the rat aorta.

Y axis represents the isotope activity in count per minute (cpm); 6-k-PGF_1α_: 6-keto prostaglandin PGF_1α_; TxB_2_: thromboxane B_2_; All data are represented as mean ± SD. n=9−9 rats/group. One-way ANOVA, Bonferroni test, *p < 0.05, **p < 0.03, ***p < 0.01 compared to the diabetic vehicle-treated groups.

The production of CON metabolites of the diabetic rat aorta was significantly reduced by (S)-L1 compared to the vehicle-treated diabetic groups. The synthesis of DIL metabolites in the diabetic rat abdominal aorta was stimulated by (S)-L1 and NE-100 compared to the vehicle-treated diabetic samples. The (S)-L1 and the NE-100 ligands significantly decreased the CON/DIL ratio of the metabolites in the diabetic rat abdominal aorta ring, compared to the vehicle-treated diabetic samples *(Table 8)*.

**Table 8.**
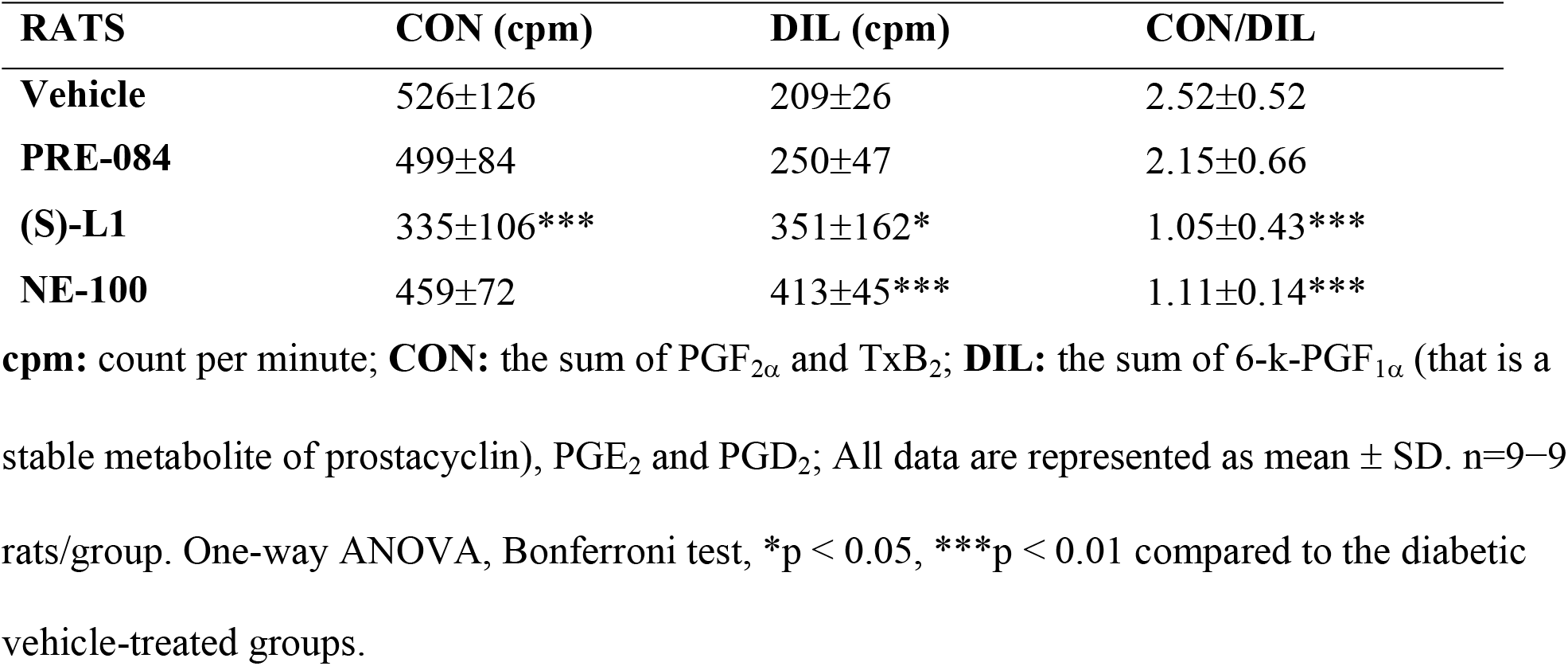
Vasoconstrictor, platelet aggregator (CON) and vasodilator, platelet anti-aggregator (DIL) COX metabolites of diabetic rat abdominal aorta.

| RATS | CON (cpm) | DIL (cpm) | CON/DIL |
| --- | --- | --- | --- |
| Vehicle | 526±126 | 209±26 | 2.52±0.52 |
| PRE-084 | 499±84 | 250±47 | 2.15±0.66 |
| (S)-L1 | 335±106*** | 351±162* | 1.05±0.43*** |
| NE-100 | 459±72 | 413±45*** | 1.11±0.14*** |

**cpm:** count per minute; **CON:** the sum of $\text{PGF}_{2\alpha}$ and $\text{TxB}_2$ ; **DIL:** the sum of 6-k- $\text{PGF}_{1\alpha}$ (that is a stable metabolite of prostacyclin), $\text{PGE}_2$ and $\text{PGD}_2$ ; All data are represented as mean ± SD. n=9–9 rats/group. One-way ANOVA, Bonferroni test, \* $p < 0.05$ , \*\*\* $p < 0.01$ compared to the diabetic vehicle-treated groups.

##### COX enzymes levels in the diabetic abdominal rat aorta measured by ELISA

In the diabetic rat aorta, no change was seen except for the S1R antagonist NE-100 group, in which significantly reduced COX-1 concentration was measured compared to the vehicle-treated diabetic animals shifting the balance COX-1/COX-2 toward COX-2. *(Table 9)*.

**Table 9.** Effect of S1R ligands on the levels of COX-1 and COX-2 enzymes determined by ELISA in diabetic rat abdominal aorta.

| RATS | COX-1 | COX-2 | COX-1+COX- | COX-1/COX-2 |
| --- | --- | --- | --- | --- |
| Vehicle | 3.72±1.02 | 7.93±2.4 | 11.15±4.07 | 0.47±0.04 |
| PRE-084 | 3.93±0.27 | 7.57±0.98 | 11.97±0.78 | 0.51±0.09 |
| (S)-L1 | 3.75±1.06 | 7.99±0.52 | 12.64±0.71 | 0.51±0.05 |
| NE-100 | 2.43±1.11* | 8.07±0.97 | 10.31±1.25 | 0.40±0.24 |

COX-1: Type 1 cyclooxygenase; COX-2: Type 2 cyclooxygenase; All data are represented as mean ± SD. n=9-9 rats/group. One-way ANOVA, Bonferroni test, \*p < 0.05 compared to the diabetic vehicle-treated groups.

## 4. Discussion

In the present study, we investigated the effects of sub-chronic, in vivo administered S1R ligands on *ex vivo/in vitro* eicosanoid synthesis of STZ-induced diabetic male rat platelets and abdominal aorta. We selected STZ-induced diabetes mellitus as a reliable, reproducible model with chronic inflammation and oxidative stress in which the development and time course of complications can be easily monitored.31

### STZ-induced diabetic model

STZ contains glucose and N-methyl-N-nitrosocarbamide groups.^32^ Its glucose component binding to glucose transporter-2 (GLUT-2) promotes the entry of STZ into pancreatic, liver and renal tubule epithelial cells with GLUT-2.^33,34^ The N-methyl-N-nitrosourea constituent of STZ thus transported into cells induces DNA methylation,^35^ alkylation^36^ and oxidation.^37^ These processes inducing cell apoptosis, lead to the development of diabetes mellitus and liver and kidney damage.

In the present study, the development of STZ-induced diabetes mellitus of the rats, was supported by elevated serum glucose level (hyperglycemia) and reduced rate of body weight gain, comparing to the physiological. None of the ligands that we administered i.p. for one week at a dose of 3 mg/kg/day significantly altered the STZ-induced increase in serum glucose concentration and body retardation *(Table 1)*, i.e. they couldn't restore the STZ-induced metabolic changes.

The increase in serum cholesterol, ALT and urea levels in STZ-induced diabetic animals could be explained by the hepatic and renal toxic effects of STZ.^33,34^ All of the sigma-1 ligands we tested similarly enhanced the STZ-induced increase in serum ALT concentrations indicative of hepatotoxicity *(Table 2)*. This may be explained by the metabolism of the ligands in the liver.^38^

The platelet count below the reference value,^30^ already observed in our previous studies, could be explained by reduced thrombopoietin production due to the liver and kidney damage associated with STZ-induced diabetes.^39^ Sub-chronic treatment with S1R ligands had, no effect on platelet formation, although this is not relevant in the present study as eicosanoid synthesis was monitored at a standard platelet count (2.5×10^8^ platelets/μL) *(Table 1)*.

### Sigma-1 receptor ligands

Since the presence of S1R protected against pathological changes in a model of STZ-induced diabetes mellitus^14^ we hypothesized S1R ligands may also modify endothelial and platelet function in this model. For our studies, we selected PRE-084, NE-100 and (S)-L1 from among the S1R ligands, based on their binding strength to the receptor and their binding site in the S1R binding pocket as determined in our previous experiments.^23^ Based on platelet lifespan, the *in vivo* treatment duration was one week.^40,41^ Preliminary time-dependent studies of the serum levels of i.p. administered S1R ligands demonstrated that all ligands entered the circulation, but 20 h after the last treatment, their serum concentrations were only at the limit of detection *(Table 3)*, i.e. a direct acute effect of ligands could be excluded.^23^ Thus, the changes in platelet and abdominal aortic AA metabolism *ex vivo* can be attributed to the *in vivo* effects of S1R ligands on platelet and aortic function.

Sub-chronic, *in vivo* treatment with S1R ligands was also able to alter the AA metabolism of platelets and aorta of STZ-induced diabetic rats, despite the ligands no being present in the blood at the time of *ex vivo/in vitro* study *(Table 4 and 7)*.

S1R ligands can modulate the PL content of the cell membrane, the re- or de-acylation of PLs, as well as the amount of free AA released from PLs by phospholipases.^4,42,43^ They may also affect the expression and/or activity of COX and LOX enzymes involved in AA metabolism. From the two COX isoforms, the COX-1 constitutively expressed in various tissues, but the expression of COX-2 is mostly inducible.^44^

### Effects of S1R ligands on the level of Sigmar1 and Ptgs1 mRNA in rat platelets

Although the platelets are anucleated cells, they can *de novo* synthesize COX-1 from the cytoplasmic mRNA that originates from the megakaryocyte.^45^ In human platelets, Hu and co-workers^46^ have also detected COX-2 mRNA and protein, although at significantly lower levels than COX-1, in concordance with our data.

However, under the current experimental conditions, we could not detect *Ptgs2* mRNA in platelets from diabetic rats until cycle 40 of the RT-qPCR. This may be explained by the fact that only limited amounts of megakaryocyte mRNA transcripts are available for protein synthesis (e.g. COX-2), and therefore, in diabetes mellitus, increased *in vivo* AA metabolism may lead to depletion of intracellular reserves (mRNA, enzyme pool, [Ca^2+^]_i_).

Of the S1R ligands used *in vivo*, PRE-084 and (S)-L1 increased platelet *Sigmar1* transcript levels *(Fig. 1)*, but did not modify *Ptgs1 (Fig. 2)*. In contrast, NE-100 only induced *Ptgs1* transcript reduction *(Fig. 2)*.

### Effects of S1R ligands on the ex vivo eicosanoid synthesis of platelets

Treatment with PRE-084 and (S)-L1 ligands, although not affecting the total amount of radioactive eicosanoids (COX+LOX) synthesized by platelets from diabetic rats, shifted AA metabolism towards the LOX pathway due to a reduction in COX metabolites *(Table 4)*. These results suggest that neither PRE-084 nor (S)-L1 altered the release of free AA substrate from PLs. Since we detected unchanged *Ptgs1* mRNA *(Fig. 2)*, elevated COX-1 and COX-2 enzyme concentrations by ELISA (Table 6) and unchanged radioactive LOX product (Table 4) formation with decreased COX metabolite *(Table 4)* synthesis, this may suggest that PRE-084 and (S)-L1 have platelet COX enzyme inhibitory effects. Despite a decrease in the total amount of radioactive COX metabolites, PRE-084 and (S)-L1 ligands resulted in an increase in the CON/DIL ratio, indicating a predominance of platelet aggregating and vasoconstrictor COX metabolites *(Fig. 3, Table 5)*.

In contrast to PRE-084 and (S)-L1 ligand, NE-100 decreased platelet *Ptgs1* mRNA levels (Fig. 2), did not modulate COX-1, but increased COX-2 enzyme levels determined by ELISA *(Table 6)*. Nevertheless, it did not affect the total amount of radioactive COX metabolites (Table 4), which resulted from the combined action of COX-1 and COX-2 enzymes. NE-100 ligand did not alter the proportion of CON or DIL metabolites produced by the COX pathway *(Fig. 3 Table 5)*. Since NE-100 significantly reduced the total amount of radioactive AA metabolites (COX+LOX) but did not induce a shift in the COX/LOX ratio *(Table 4)*, its effect may be due to inhibition of phospholipase, i.e. reduction of the free AA substrate.

### Effects of S1R ligands on the ex vivo eicosanoid synthesis of abdominal aorta

The effects of all three sigma-1 ligands we studied on abdominal aortic eicosanoid synthesis differed widely. The total amount of radioactive AA products (COX+LOX) in the aorta was not significantly affected by PRE-084, decreased by (S)-L1, whereas enhanced by NE-100. However, since neither (S)-L1 nor NE-100 resulted in a COX/LOX ratio shift, (S)-L1 may have reduced, whereas NE-100 may have enhanced, aortic free AA release. In contrast to platelets, PRE-084 and (S)-L1 ligands did not alter the total amount of aortic radioactive COX metabolites *(Table 7)*, as confirmed by determination of COX concentration by ELISA *(Table 9)*, but they did decrease the formation of LOX products *(Table 7)*. Regardless of the total amount of radioactive COX metabolites, both (S)-L1 and NE-100 resulted in a decrease in the CON/DIL ratio, with a consequent predominance of the synthesis of vasodilator eicosanoids that inhibit platelet aggregation *(Table 8)*. In other words, sigma-1 ligands are also able to modulate specific enzymes involved in the synthesis of certain eicosanoids.

Under physiological conditions, it is known that platelet and endothelial cell AA metabolism differ significantly. Platelets primarily produce the vasoconstrictor thromboxane, which induces platelet aggregation, and the endothelium primarily produces the vasodilator prostacyclin, which inhibits platelet aggregation, but under physiological conditions these products are in equilibrium to ensure normal local circulation.^31,39,44^

Taken together, ligands have opposite effects on platelet and aortic AA metabolism in the diabetic rat, which may promote the pursuit of physiological equilibrium, even under pathological conditions.

## 5. Conclusion

Sub-chronic *in vivo* treatment of STZ-induced diabetic rats with PRE-084, (S)-L1, NE-100 S1R ligands modulated both platelet and aortic *ex vivo* eicosanoid synthesis.

The new (S)-L1 ligand, STZ-induced diabetic platelet and aortic eicosanoid synthesis was similar to, although different in magnitude from, the S1R agonist PRE-084 in most of the parameters we studied, whereas the S1R antagonist NE-100 ligand had opposite effects. Our results suggest that S1R ligands may play a role in the regulation of cellular functions and local circulation by affecting AA metabolism (transcription, translation, enzyme induction). In certain pathological conditions, these ligand and cell-specific effects may play a compensatory role, i.e. they may help to restore physiological balance.

## Author Contributions

Conceptualization, S.V. and Z.M.; methodology, S.V., L.B., K.L., F.T; formal analysis, S.V., L.B. Z.M.; investigation, S.V., L.B., F.T.; resources, S.V., B.P., T.J., M.A.D., Z.M.; data curation, S.V., L.B., K.L., F.T., G.R., Z.M.; writing—original draft preparation, S.V., Z.M.; writing—review and editing, L.B., K.L., G.R., B.P., L.F., T.J., M.A.D; visualization, S.V., L.B., F.T., Z.M.; supervision, G.R., B.P., L.F., T.J., M.A.D., Z.M.; funding acquisition, S.V., B.P., T.J., M.A.D., Z.M.. All authors have read and agreed to the published version of the manuscript.”

## Institutional Review Board Statement

Animal experiments were performed under the protocol accepted by the Ethical Committee for the Protection of Animals in Research at the University of Szeged, Hungary (Permit No. X./238/2019.). All experiments were carried out in accordance with the Guide for the Care and Use of Laboratory Animals published by the US National Institutes of Health.

## Data Availability Statement

The data that support the findings of this study are available from the corresponding author upon reasonable request.

## Acknowledgments

We are grateful to the staff of the Department of Laboratory Medicine, Albert Szent-Györgyi Medical School, University of Szeged, for the determination of the laboratory parameters reported in this communication.

## References

1. American Diabetes Association. Diagnosis and classification of diabetes mellitus. Diabetes Care 2014; 37 Suppl 1: S81–90.

2. Carrizzo A, Izzo C, Oliveti M, et al. The Main Determinants of Diabetes Mellitus Vascular Complications: Endothelial Dysfunction and Platelet Hyperaggregation. Int J Mol Sci; 19. Epub ahead of print 28 September 2018. DOI: 10.3390/ijms19102968.

3. Poznyak A, Grechko AV, Poggio P, et al. The Diabetes Mellitus-Atherosclerosis Connection: The Role of Lipid and Glucose Metabolism and Chronic Inflammation. Int J Mol Sci 2020; 21: E1835.

4. Paes AM de A, Gaspar RS, Fuentes E, et al. Lipid Metabolism and Signaling in Platelet Function. Adv Exp Med Biol 2019; 1127: 97–115.

5. Su T-P, Su T-C, Nakamura Y, et al. The Sigma-1 Receptor as a Pluripotent Modulator in Living Systems. Trends Pharmacol Sci 2016; 37: 262–278.

6. Aishwarya R, Abdullah CS, Morshed M, et al. Sigmar1’s Molecular, Cellular, and Biological Functions in Regulating Cellular Pathophysiology. Front Physiol 2021; 12: 705575.

7. Wang J, Xu D, Shen L, et al. Anti-inflammatory and analgesic actions of bufotenine through inhibiting lipid metabolism pathway. Biomed Pharmacother 2021; 140: 111749.

8. Amer MS, McKeown L, Tumova S, et al. Inhibition of endothelial cell Ca2+ entry and transient receptor potential channels by Sigma-1 receptor ligands. Br J Pharmacol 2013; 168: 1445–1455.

9. Váczi S, Barna L, Harazin A, et al. S1R agonist modulates rat platelet eicosanoid synthesis and aggregation. Platelets. Epub ahead of print 2021. DOI: 10.1080/09537104.2021.1981843.

10. Rosen DA, Seki SM, Fernández-Castañeda A, et al. Modulation of the sigma-1 receptor-IRE1 pathway is beneficial in preclinical models of inflammation and sepsis. Sci Transl Med 2019; 11: eaau5266.

11. Gao Q-J, Yang B, Chen J, et al. Sigma-1 Receptor Stimulation with PRE-084 Ameliorates Myocardial Ischemia-Reperfusion Injury in Rats. Chin Med J (Engl) 2018; 131: 539–543.

12. Nardai S, László M, Szabó A, et al. N,N-dimethyltryptamine reduces infarct size and improves functional recovery following transient focal brain ischemia in rats. Exp Neurol 2020; 327: 113245.

13. Tagashira H, Matsumoto T, Taguchi K, et al. Vascular endothelial σ1-receptor stimulation with SA4503 rescues aortic relaxation via Akt/eNOS signaling in ovariectomized rats with aortic banding. Circ J 2013; 77: 2831–2840.

14. Ha Y, Saul A, Tawfik A, et al. Diabetes accelerates retinal ganglion cell dysfunction in mice lacking sigma receptor 1. Mol Vis 2012; 18: 2860–2870.

15. Su TP, Wu XZ, Cone EJ, et al. Sigma compounds derived from phencyclidine: identification of PRE-084, a new, selective sigma ligand. J Pharmacol Exp Ther 1991; 259: 543–550.

16. Allahtavakoli M, Jarrott B. Sigma-1 receptor ligand PRE-084 reduced infarct volume, neurological deficits, pro-inflammatory cytokines and enhanced anti-inflammatory cytokines after embolic stroke in rats. Brain Res Bull 2011; 85: 219–224.

17. Dong H, Ma Y, Ren Z, et al. Sigma-1 Receptor Modulates Neuroinflammation After Traumatic Brain Injury. Cell Mol Neurobiol 2016; 36: 639–645.

18. Motawe ZY, Abdelmaboud SS, Cuevas J, et al. PRE-084 as a tool to uncover potential therapeutic applications for selective sigma-1 receptor activation. Int J Biochem Cell Biol 2020; 126: 105803.

19. Okuyama S, Nakazato A. NE-100: A Novel Sigma Receptor Antagonist. CNS Drug Reviews 1996; 2: 226–237.

20. Dvorácskó S, Lázár L, Fülöp F, et al. Novel High Affinity Sigma-1 Receptor Ligands from Minimal Ensemble Docking-Based Virtual Screening. Int J Mol Sci 2021; 22: 8112.

21. Furman BL. Streptozotocin-Induced Diabetic Models in Mice and Rats. Curr Protoc Pharmacol 2015; 70: 5.47.1–5.47.20.

22. Trufelli H, Palma P, Famiglini G, et al. An overview of matrix effects in liquid chromatography-mass spectrometry. Mass Spectrom Rev 2011; 30: 491–509.

23. Váczi S, Barna L, Laczi, Krisztián, et al. Effects of sub-chronic, in vivo administration of sigma-1 receptor ligands on platelet and aortic arachidonate cascade in rat. Eur J Pharmacol. Under Review

24. Mezei Z, Kis B, Gecse A, et al. Platelet eicosanoids and the effect of captopril in blood pressure regulation. Eur J Pharmacol 1997; 340: 67–73.

25. Mezei Z, Kis B, Gecse A, et al. Platelet arachidonate cascade of migraineurs in the interictal phase. Platelets 2000; 11: 222–225.

26. Mezei Z, Zamani-Forooshani O, Csabafi K, et al. The effect of kisspeptin on the regulation of vascular tone. Can J Physiol Pharmacol 2015; 93: 787–791.

27. Abdel-Halim MS, Lundén I, Cseh G, et al. Prostaglandin profiles in nervous tissue and blood vessels of the brain of various animals. Prostaglandins 1980; 19: 249–258.

28. Kis B, Szabó CA, Pataricza J, et al. Vasoactive substances produced by cultured rat brain endothelial cells. Eur J Pharmacol 1999; 368: 35–42.

29. Cryer B. Management of patients with high gastrointestinal risk on antiplatelet therapy. Gastroenterol Clin North Am 2009; 38: 289–303.

30. Charles R. Clinical Laboratory Parameters for Crl:WI(Han) Rats, https://www.criver.com/sites/default/files/Technical%20Resources/Clinical%20Laboratory%20Parameters%20for%20Crl-WI(Han)%20Rats%20-%20March%202008.pdf (2008).

31. Csanyi G, Lepran I, Flesch T, et al. Lack of endothelium-derived hyperpolarizing factor (EDHF) up-regulation in endothelial dysfunction in aorta in diabetic rats. Pharmacol Rep 2007; 59: 447–455.

32. Elsner M, Guldbakke B, Tiedge M, et al. Relative importance of transport and alkylation for pancreatic beta-cell toxicity of streptozotocin. Diabetologia 2000; 43: 1528–1533.

33. Eleazu CO, Iroaganachi M, Eleazu KC. Ameliorative Potentials of Cocoyam (*Colocasia esculenta* L.) and Unripe Plantain (*Musa paradisiaca* L.) on the Relative Tissue Weights of Streptozotocin-Induced Diabetic Rats. Journal of Diabetes Research 2013; 2013: 1–8.

34. Valentovic MA, Alejandro N, Betts Carpenter A, et al. Streptozotocin (STZ) diabetes enhances benzo(alpha)pyrene induced renal injury in Sprague Dawley rats. Toxicol Lett 2006; 164: 214–220.

35. Szkudelski T. Streptozotocin–nicotinamide-induced diabetes in the rat. Characteristics of the experimental model. Exp Biol Med (Maywood) 2012; 237: 481–490.

36. Friederich M, Hansell P, Palm F. Diabetes, Oxidative Stress, Nitric Oxide and Mitochondria Function. CDR 2009; 5: 120–144.

37. Gul M, Laaksonen DE, Atalay M, et al. Effects of endurance training on tissue glutathione homeostasis and lipid peroxidation in streptozotocin-induced diabetic rats: Training and glutathione in diabetic rats. Scandinavian Journal of Medicine & Science in Sports 2002; 12: 163–170.

38. Yamamoto T, Hagima N, Nakamura M, et al. Prediction of differences in *in vivo* oral clearance of *N*, *N*-dipropyl-2-[4-methoxy-3-(2-phenylethoxy)phenyl]ethylamine monohydrochloride (NE-100) between extensive and poor metabolizers from *in vitro* metabolic data in human liver microsomes lacking CYP2D6 activity and recombinant CYPs. Xenobiotica 2004; 34: 687–703.

39. Mezei Z, Váczi S, Török V, et al. Effects of kisspeptin on diabetic rat platelets. Can J Physiol Pharmacol 2017; 95: 1319–1326.

40. Thon JN, Italiano JE. Platelets: production, morphology and ultrastructure. Handb Exp Pharmacol 2012; 3–22.

41. Cimmino G, Golino P. Platelet biology and receptor pathways. J Cardiovasc Transl Res 2013; 6: 299–309.

42. Starr JB, Werling LL. Sigma-receptor regulation of [3H]arachidonic acid release from rat neonatal cerebellar granule cells in culture. J Neurochem 1994; 63: 1311–1318.

43. Hayashi T. The Sigma-1 Receptor in Cellular Stress Signaling. Front Neurosci 2019; 13: 733.

44. Mitchell JA, Kirkby NS. Eicosanoids, prostacyclin and cyclooxygenase in the cardiovascular system. Br J Pharmacol 2019; 176: 1038–1050.

45. Evangelista V, Manarini S, Di Santo A, et al. De novo synthesis of cyclooxygenase-1 counteracts the suppression of platelet thromboxane biosynthesis by aspirin. Circ Res 2006; 98: 593–595.

46. Hu Q, Cho MS, Thiagarajan P, et al. A small amount of cyclooxygenase 2 (COX2) is constitutively expressed in platelets. Platelets 2017; 28: 99–102.

